# Unique rhythmic signatures in arrhythmic birdsong

**DOI:** 10.64898/2025.12.19.695402

**Authors:** Anna N Osiecka, Javier Oñate-Casado, Tereza Petrusková, Adrián Barrero, Václav Beran, Tereza Kubíková, Cristian Pérez-Granados, Michal Porteš, Juan Traba, Adam Petrusek, Lara S Burchardt

## Abstract

Animal rhythms are gaining increasing attention in the studies of behaviour and musicology. Recently, it has been shown that rhythm itself can be used as an information coding channel. Now we ask: does this hold true for arrhythmic sequences? To answer that, we analysed songs of the tawny pipits (*Anthus campestris*), migratory songbirds known for their simple songs. Using focal recordings of 384 individuals from six populations collected across their European breeding range, we calculated an extensive set of rhythmic and temporal indices to describe each song. First, the pattern of these songs was inspected and shown to be arrhythmic. Second, songs belonging to specific individuals and populations were compared using permuted discriminant function analysis and supervised uniform manifold approximation and projection. To describe the level of individuality of the rhythmic structure alone, we calculated Beecher’s statistic for all songs, as well as the potential of identity coding for each parameter separately. We show that tawny pipit males sing using individual rhythmic patterns in their arrhythmic songs, and that modern rhythmic indices, such as beat precision and integer ratios, are among some of the most individually distinct parameters of their songs. Furthermore, contrary to previous investigations based on the spectral shape and basic frequency and temporal characteristics of these songs, we show that the species displays population-specific temporal patterns, with significant differences throughout its European range. This study is the first to demonstrate geographic scale differences in birdsong rhythm, and to show that rhythmic analysis can provide useful descriptions of arrhythmic sequences.

## Introduction

Animal sounds, and birdsong in particular, have always accompanied and fascinated humans. Within the Western musical tradition alone, written proof of avian-inspired music reaches at least the 13th century (such as the Wessex *Sumer is icumen in* rota following cuckoo calls, with a preserved manuscript from 1261-1264), and is still with us in the TikTok era and Sy Wylie’s Bird Beats. In fact, some of the early attempts to describe avian sounds rely on mimicking their temporal patterns, including mnemonic phrases to recognise species (e.g. *cilp-calp* for the common chiffchaff, *Phylloscopus collybita*, in Polish and Czech, or *a little bit of bread and no cheese, please* for the yellowhammer, *Emberiza citrinella*, in English). Simple music notation attempts also tried to capture the calls’ or songs’ overall structure - these range from Athanasius Kircher’s 1650 *Musurgia Universalis* musical drawings of hens to Friederich Goethe’s *Möwenmusik* (seagull music), showcasing the seabird calls of the German Baltic coast [1].

These long-standing attempts to capture avian temporal structure underscore avian sounds not only as spectral objects but as patterned sequences in time. Nonetheless, with the development of modern signal analysis methods, quantitative analyses in bioacoustics have historically privileged spectral descriptors, leaving rhythmic organisation comparatively underexplored. Rhythm is now coming back into focus in the studies of animal behaviour thanks to the development of new, robust rhythm analysis methods [2] [3] [4]. Modern rhythmic indices provide sequence-level descriptors that quantify relative timing between elements, going beyond measures based solely on mean durations, tempos, or inter-onset intervals.

Recently, we have shown that modern rhythmic indices [4] (as opposed to simple descriptions of the temporal structure) can not only describe a sequence, but prove reliable and important sources of information for isochronous (equally distributed in time [5]) and *rallentando* (slowing down [6]) vocal sequences. Can the same methods be used for signals that appear irregular or unpredictable? This distinction is crucial because many animal displays lack overt periodicity, yet exhibit consistent internal temporal relationships that may encode relevant information (such as caller’s identity in the seemingly arrhythmic vocal displays of the tambourine dove, *Turtur tympanistria* [7]). A question thus arises, whether modern rhythmic indices can provide useful information for arrhythmic sequences? In other words, if we approach rhythm as a description of patterned temporal relationships of all elements in a sequence to each other [4], can these relationships be used to describe constructs without repeated, predictable patterns?

Testing this requires species whose songs are simple, variable, and rhythmically unpatterned. Tawny pipits (*Anthus campestris*) provide an ideal system: males produce short, simple territorial songs that vary substantially among individuals [8], [9] yet lack obvious repeated temporal patterns (Figure 1). Additionally, with substantial element overlaps, tawny pipit song presents an interesting question as to what constitutes a unit of the temporal structure.

**Figure 1:**
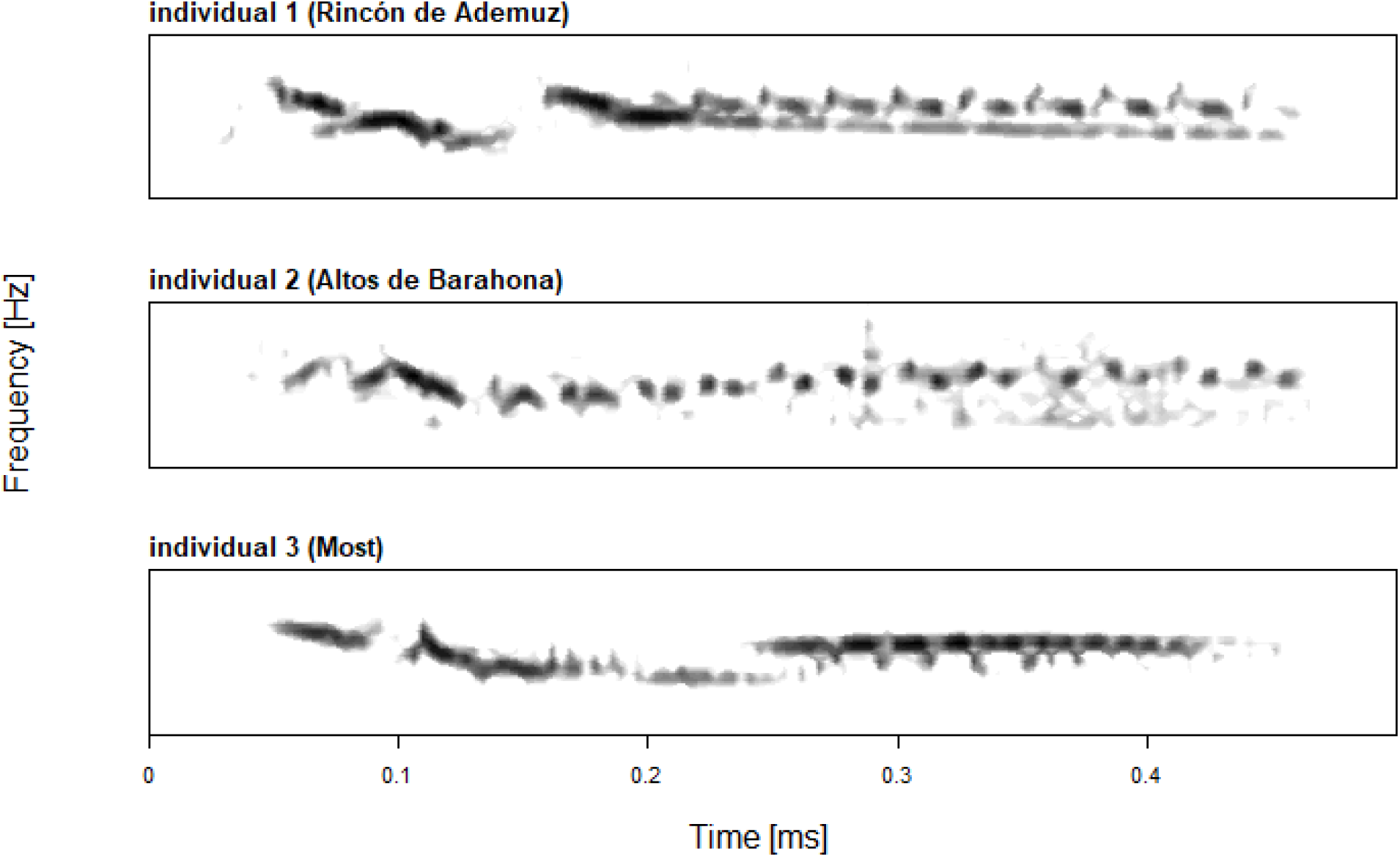
Sample songs of three different tawny pipit singers. Overall, the songs of the species are simple, often considered as one syllable, without any apparent repeated patterns. Note that while some sub-units appear to be organised and repeated (e.g. individuals 1 and 2, final sections), this is not true for all songs (individual 3). Spectra plotted using function spectro, seewave package [10], Hanning, window length 150.

This study explores the rhythmic structure of tawny pipits’ apparently arrhythmic songs as an information channel for singer identity and population of origin, by revisiting a vast recording library collected across their European breeding range (Figure 2 [11]).

**Figure 2:**
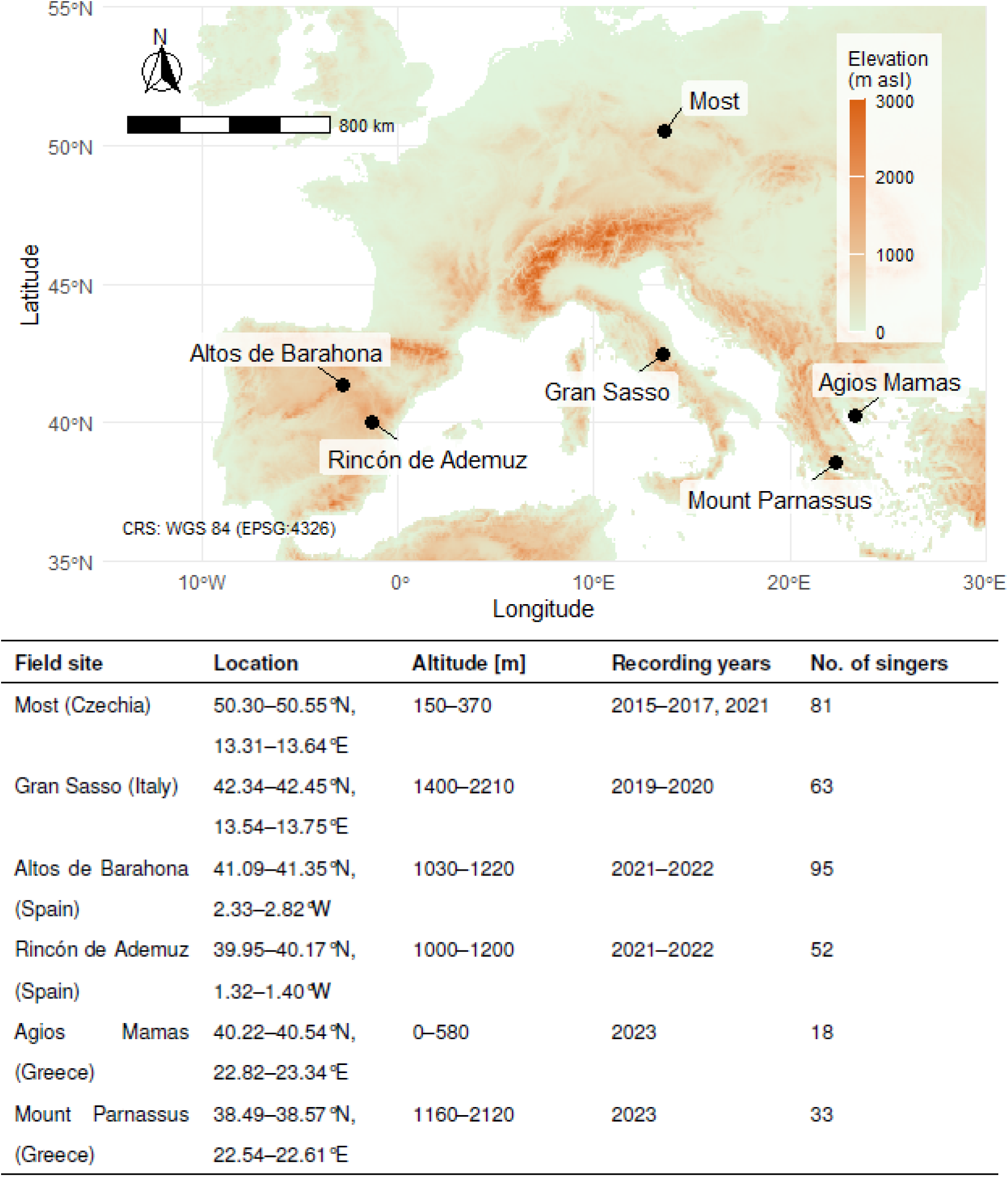
Study locations covered the species’ breeding grounds in Southern and Central Europe. Raster data obtained via *geodata* package [12]. Table presents fieldsite details and numbers of individual males recorded.

## Results

Luscinia software [13] allows users to annotate recording by "painting" directly on the spectrogram with a brush tool, tracing the spectral contours of each element. In this way, we could define the start and end times of each element by starting and stopping the brush in accordance with the visible structure. Since tawny pipit songs are not composed of clear syllables, and contain significant temporal overlaps between elements (Figure 1), we were thus able to obtain start and end points of the different songs’ elements (Figure 3) and calculate the rhythmic indices for all pairs of element onset times, independently of this overlap.

**Figure 3:**
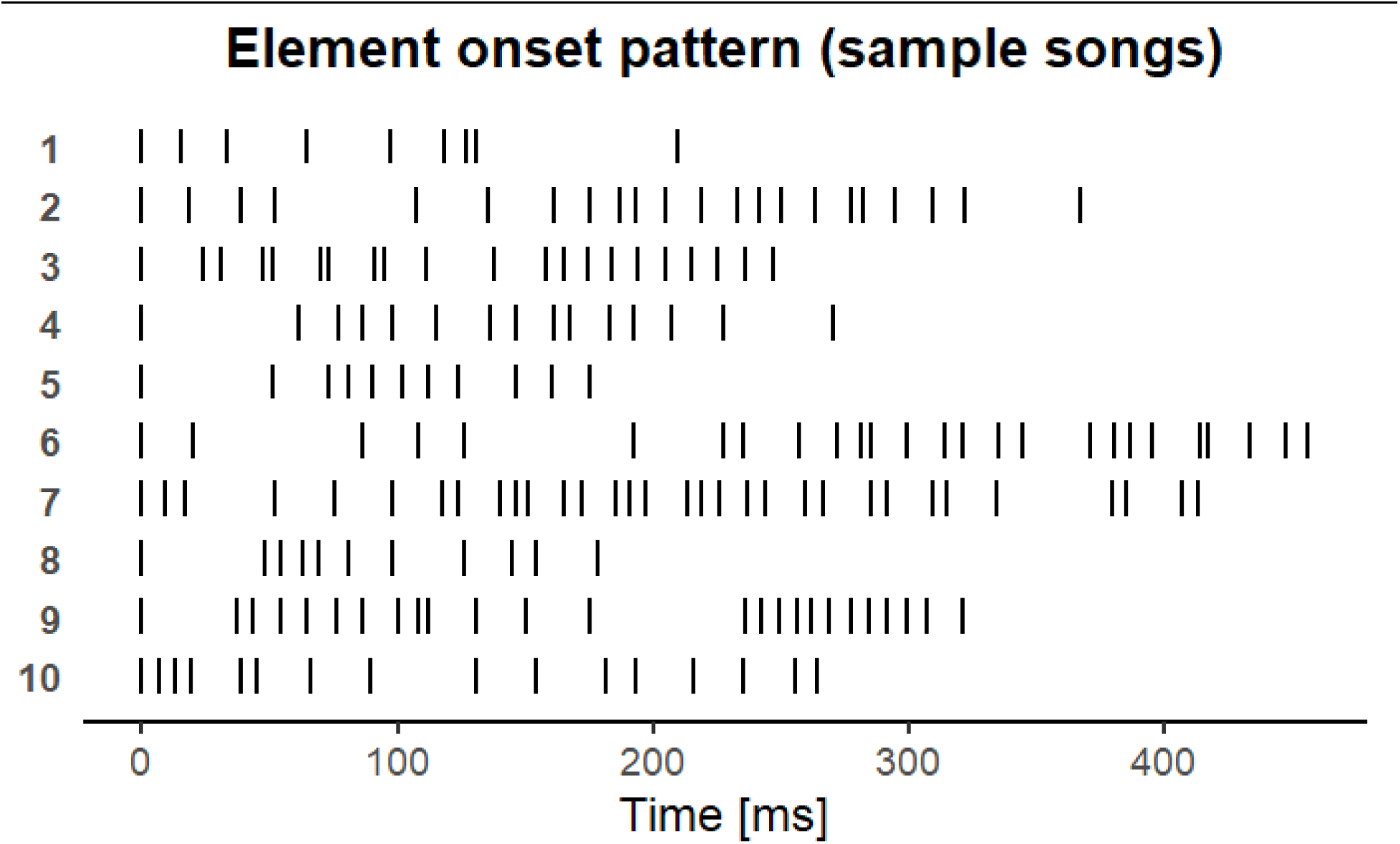
Element onset times in randomly selected sample songs of 10 random individuals indicate an apparent lack of species-specific temporal structure to tawny pipit songs

Overall, no apparent rhythmic structure emerges in the species’ song - that is, no repeated, predictable patterns emerge across the population (Figure 3). This was confirmed by plotting the distribution of integer ratios (rk) calculated for tawny pipit songs together with randomly generated integer ratios (Figure 4, panels c-d). Integer ratios represent the relationship between each two elements (e.g. ratio of durations or interonset-intervals, IOI), and are calculated for both adjacent and all element pairs in the sequence (consult Table 2 for rhythm terms and definitions).

**Figure 4:**
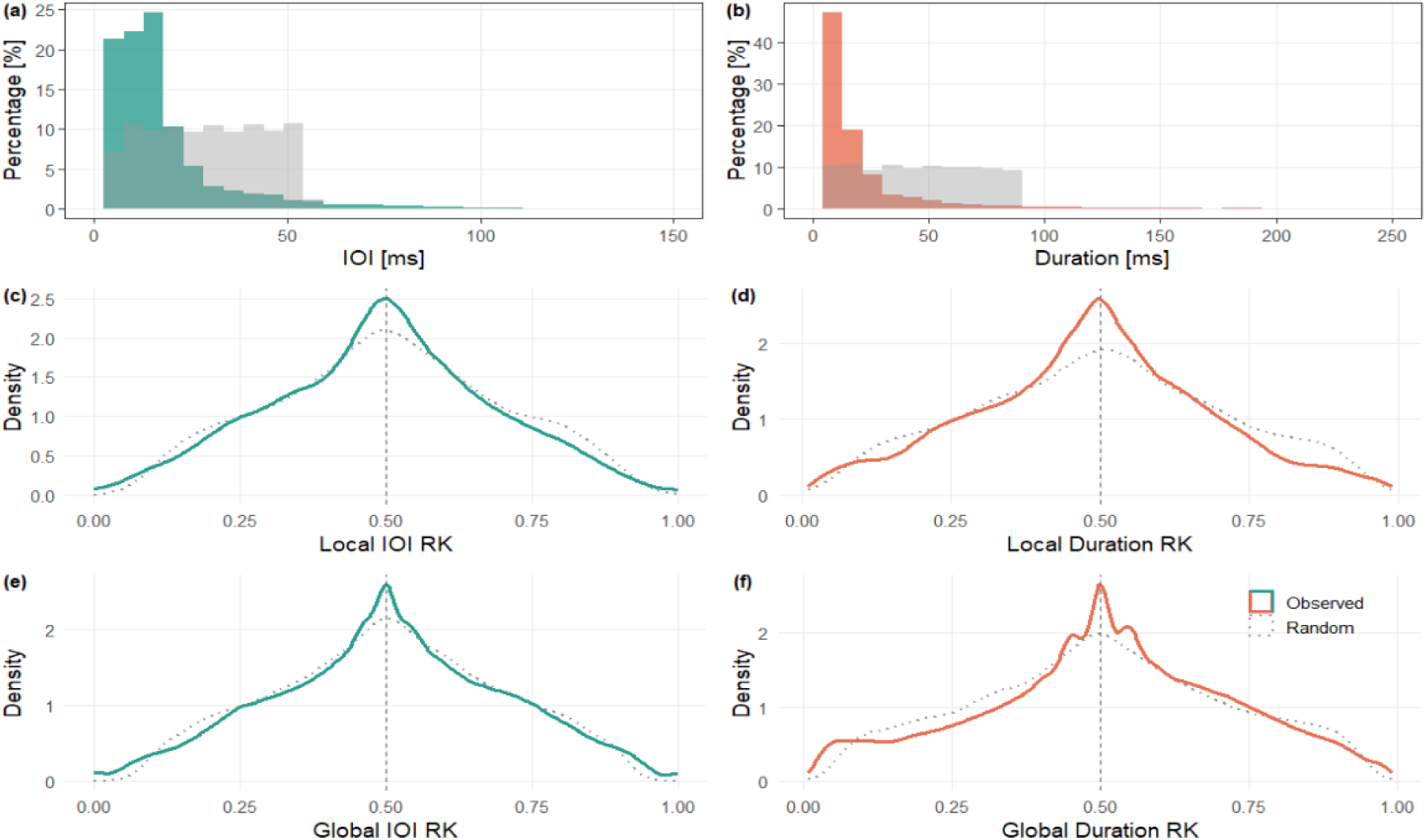
Distributions of inter-onset-intervals (panel 1), durations (panel b), and rhythmic ratios plotted with random integer ratio (rk) values (created within the 5-95% confidence interval of the original inter-onset-intervals, IOI, and durations, panels c-f). Clear peaks around a certain rk value represent the rhythmic organisation of a sequence (or, like here, a lack of thereof). Solid lines represent observed, and dotted/gray randomly generated values, green - IOI, red - duration. Indices based on observed values largely follow the distribution of these based on randomly generated ones (c-f), indicating clearly random patterns with some sub-organisation into semi- isochronous segments (0.5 density peaks within otherwise random distribution).

A closer investigation of the distributions seems to indicate smaller sub-units organised in local isochronous pattern (0.5 density peaks, Figure 4, panels c-f). This aligns with what can be observed in at least some individual songs (see e.g. individuals 1 and 2, Figure 1, as well as e.g. repeated three-element sections around 300 ms in songs 2 and 3 of individual 1, Figure 5). In other words, while there is no global rhythmic pattern for the species, individual patterns clearly emerge.

**Figure 5:**
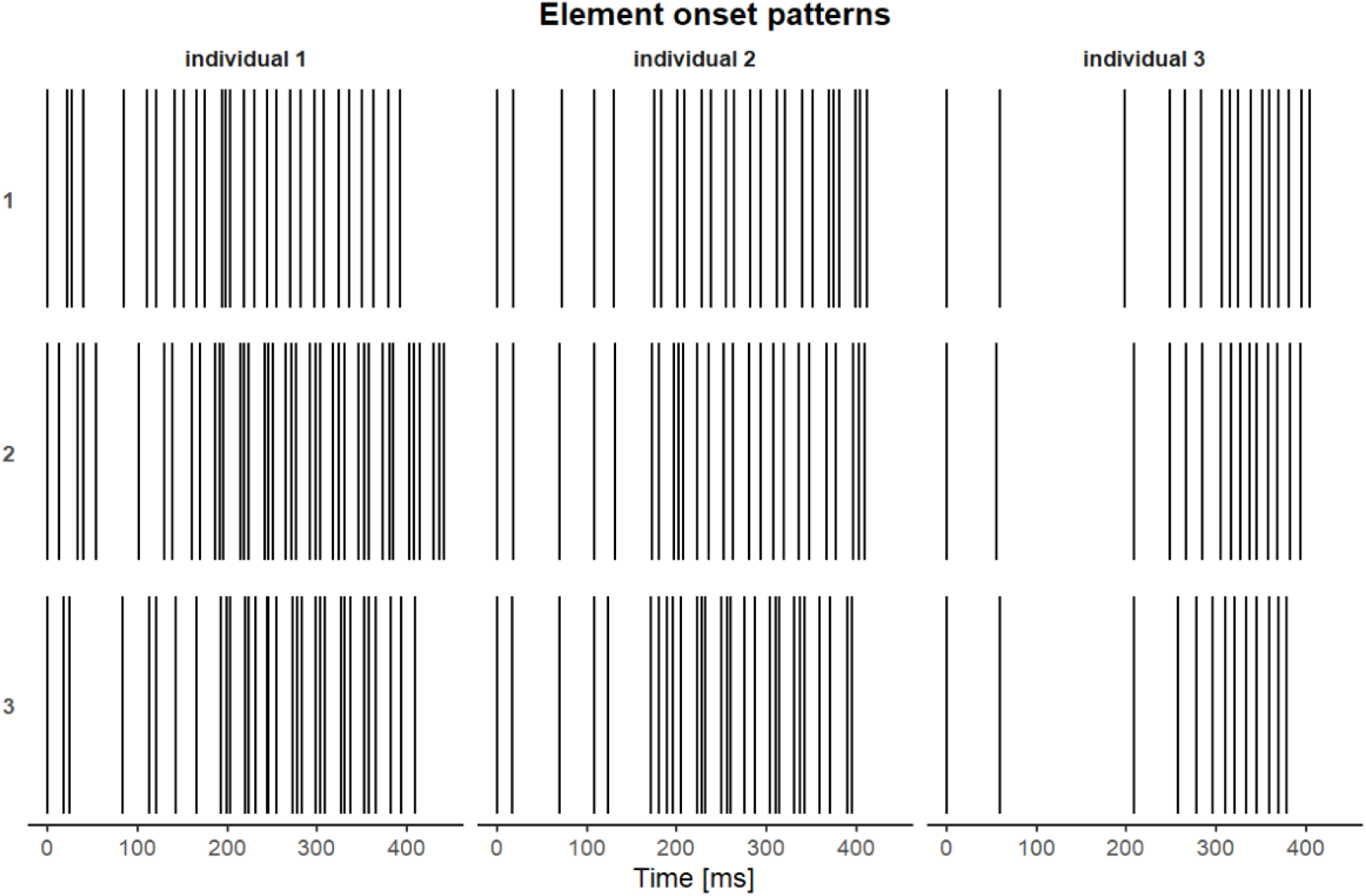
Element onset patterns for three randomly selected songs (rows) of three randomly selected individual singers (columns). First song of each individual corresponds to the spectrograms in Figure 1.

In fact, permuted discriminant function analysis (pDFA; a robust cross-classification method allowing to control for repeated measures of the same individuals and other factors potentially contributing to the songs’ structure based on the rhythmic and temporal features [14]) allowed us to classify songs to singers as well as their populations of origin with strong statistical significance, although the pDFA classification performed better at an individual scale (Table 1).

**Table 1:**
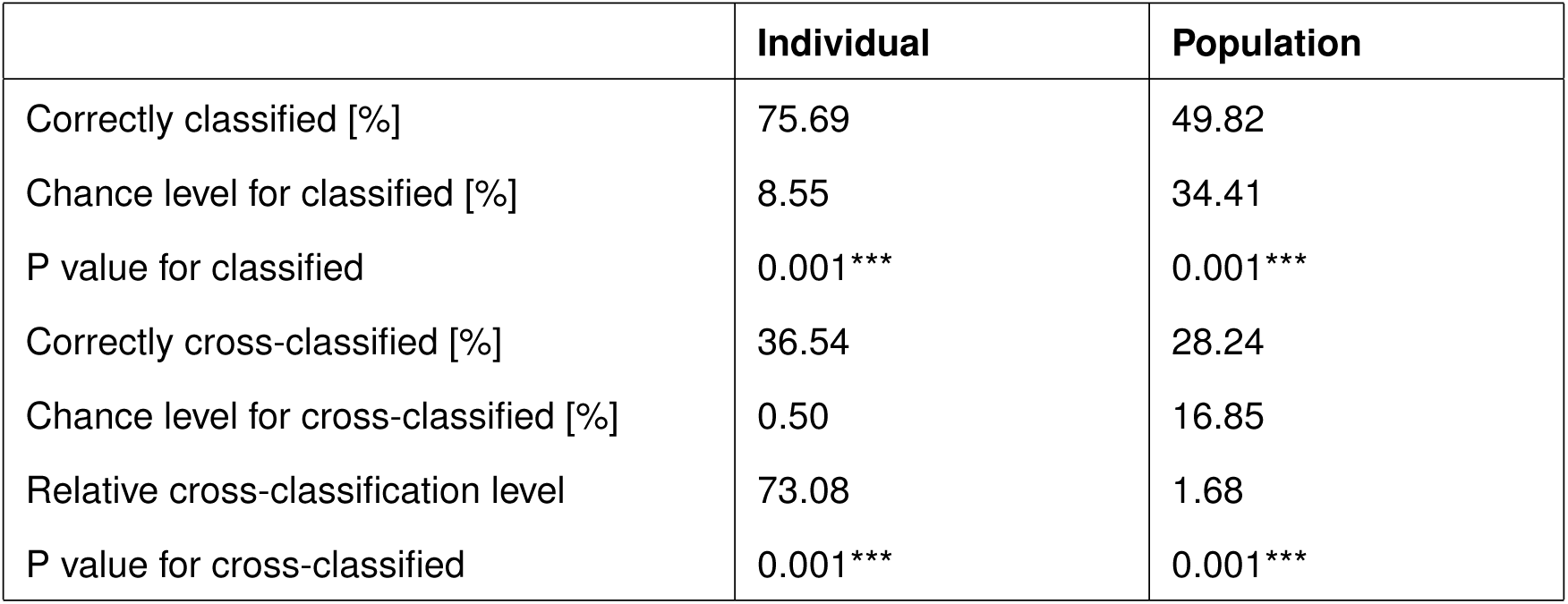
Results of the permuted discriminant function analysis classifying each song to the singer across European populations. The relative cross-classification level indicates the ratio of correct cross-classification to its chance level. Both models are based on 384 individuals, 1191 songs, 6 populations.

Supervised Uniform Manifold Aproximation and Projection (S-UMAP) clustering indicated that while some populations are clearly differentiated, others partially overlap in the abstract acoustic space (Figure 6). Basic plots of feature distribution by population can be explored in the online supplementary materials, folder plots: https://osf.io/h2fpm/overview?view_only=60916d5355de494295a1e9612ca3d40c

**Figure 6:**
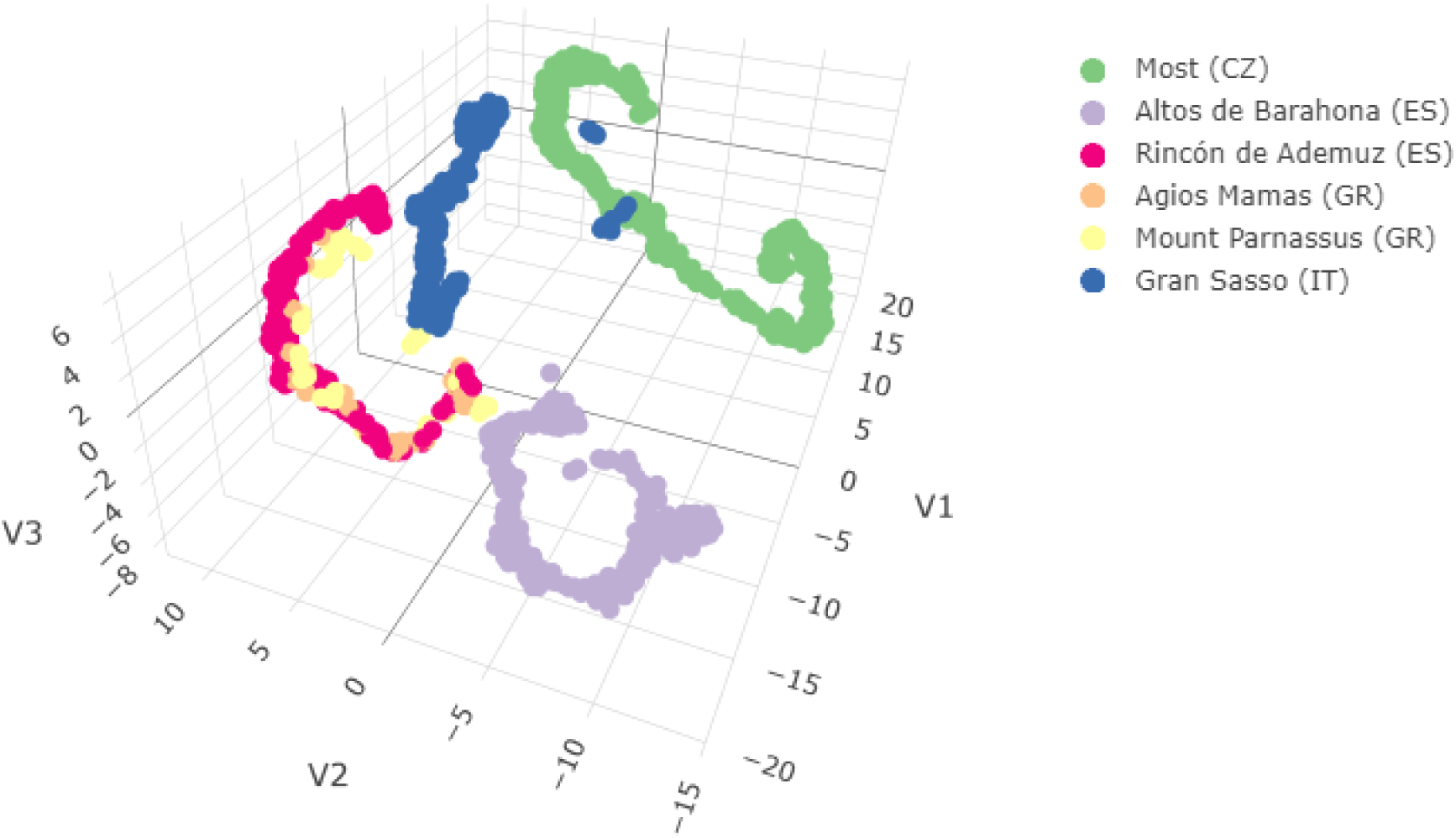
S-UMAP reveals how each song clusters in the abstract acoustic space (multidimensional sounds mapped based on their internal topology) to the singer’s population of origin (in colour). See the interactive clickable plot in the online supplementary materials: https://osf.io/h2fpm/overview?view_only=60916d5355de494295a1e9612ca3d40c)

Based on the rhythmic indices, two identity metrics commonly used as best practice in individuality studies ([15]) were calculated, namely the Beecher’s information statistic (Hs, [16]) - an indicator of the overall individuality level in a chosen population, and the potential of identity coding (PIC, 17) - which measures the contribution of each parameter to the signal’s individual traits. Hs reached 18.3 (for both *all* parameters and only these *significantly different between individuals*). All temporal and rhythmic parameters indicated strong importance for identity coding (i.e., PIC over 1, [17]), where the coefficient of variation of beat precision was prominently the most important parameter, with the PIC value of 33.36 exceeding all others by an order of magnitude (Figure 7).

**Figure 7:**
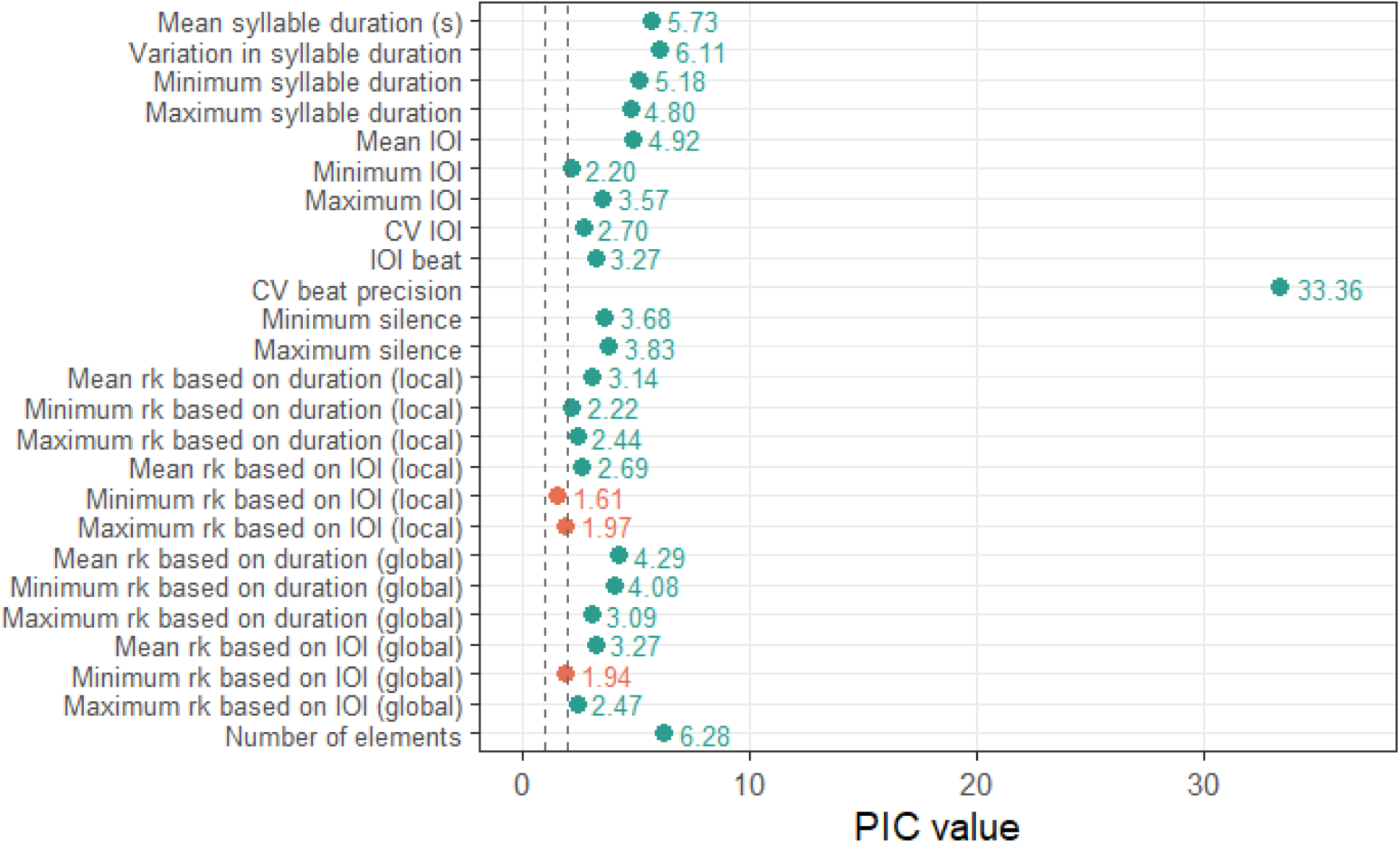
Potential of Identity Coding of the calculated temporal parameters. Dotted lines mark standard significance thresholds > 1 (red: important for identity coding) and > 2 (green: very important for identity coding). Note that each calculation is performed separately, and thus feature correlation is inconsequential here.

## Discussion

This is the first time that *rhythmic indices* have been shown useful in describing arrhythmic structures, and a source of information within them. Here, the rhythmic structure of the tawny pipits’ arrhythmic songs serve as channels carrying information on both the singer’s identity and their population of origin.

### Rhythm as a descriptor of arrhythmic sequences

Rhythm has been previously suggested [18], but to our knowledge, not used as an information coding channel in animal communication studies. Our own work used rhythmic descriptors as information sources in isochronous [5] and *rallentando* [6] vocal displays, where deviations from the perfectly isochronous beat could be expected to matter. Humans tend to find rhythm even when there is none [19] - which brings us to the assumption that the organisation of arrhythmic sequences is not meaningless The fact that rhythmic methods could be successfully used to describe arrhythmic sequences is promising in that it allows us to use them in comparative, large scale studies, and e.g. look into the evolution of rhythm or vocal (and other) signals independently of the reliability of their temporal patterning, and contribute to our understanding of the evolution of language and music [20] [21] [22] [23].

Some previous work has indicated rhythm as a possible information channel while using other methodologies. Notably, corn crake (*Crex crex*) call sequences show a somewhat syntactic modification to meaning, dependent on what appears to be integer ratios (and was tested as unquantified "relative patterning" of the calls [24]. Previous studies have also found individual and/or sex-specific patterns in simpler temporal descriptors in a wide range of taxa, such as the call or pulse rate (e.g. Cape gannet, *Morus capensis* [25], or elephant seals, *Mirounga leonina* [26]), exact note durations (e.g. tambourine dove [7], or spectral tarsiers, *Tarsius spectrumgurskyae* [27]), or exact [7] and mean syllable inter-onset-intervals (e.g. indris, *Indri indri* [28], or the little auk, *Alle alle* [29]). Even very simple temporal parameters can matter, of course - for example, call duration is extremely important in emotional communication [30]. The difference between simple temporal descriptors (also used in the present study) and rhythmic indices that capture the relative *patterning* of elements should be made clear here.

Because rhythm is often treated as an intuitively obvious concept, the term is still often misused. This highlights the importance of relying on existing definitions and approaches and refining rather than reinventing them. Over the past decade, substantial progress has been made toward developing robust methodologies, clear terminology, and comparative frameworks for rhythm studies, with best practices outlined in several key texts (e.g. [31] [32] [3] [4]).

### (Ar)rhythmic individuality

We could correctly assign the songs to the individual singer at a level high above chance, with the relative cross-classification level reaching 73.1%. While this is the first attempt focusing solely on temporal patterns, this individuality is not surprising: tawny pipit males use stereotyped songs [8], and assignment to song type is the most reliable method of identity tracking in the field recordings [9].

What stands out is the very strong contribution of the coefficient of variation of beat precision to vocal identity, reaching an outstanding PIC over 33. It should be noted that PIC values are measures of variation, and as such do not have a maximum possible value. However, surprised at this result, we have ran an additional PIC calculation based on the selected singers with the most consistent beat precision values (with coefficient of variation below 0.7; note that this is representative for the species, with most values in the range 0.65-0.70, and all over 0.64), to see whether this might be an artifact of a few very variable birds. This exclusion did not change the overall result, as the PIC value (26.8) for CV beat precision remained outstandingly high, and we decided to keep the result based on the full data in this work. In fact, variation in rhythmic and temporal parameters seems to emerge as the most important indicator of individual features in some species (e.g. age in the Cape fur seal, *Arctocephalus pusillus* [33] or identity in the little auk (PIC for CV of syllable duration equal 17.2 [29]) and herein).

Here, we are looking at a "global individuality", that is, the overall differences between birds at the European breeding grounds, but it should be noted that songbirds are known for their local variation (e.g. [34] [35] [36] [37]). In other words, within a local population one may observe yet other patterns, possibly driven by the social structure or distance between neighbouring singers (such as the apparently locally driven rhythmic diversification from the nearest singers in the yellowhammer [5]).

### Geographical variation and segregation of information

Previous work based on spectral shapes and measurements of tawny pipit songs suggested that the individuality in fact obscures geographic variation in the species [11]. Here, revisiting the same dataset from a purely rhythmic perspective, we show that geographical patterns do in fact exist in the species’ song, even though they might not be present in its spectral shape [11].

This suggests that larger-scale song evolution in tawny pipits may take place primarily in the temporal dimension, while frequency related adjustments may function at a more local scale to diversify acoustic signals among individuals within populations. At what scales exactly this segregation might happen, however, remains unclear. In the common chiffchaff, simple temporal parameters such as the number of syllables and silence durations are some of the most prominent indicators of geographical song differences, and show a progressive eastward change across the Palearctic [38]. Examining more species over their geographic range would help us understand whether these large-scale changes in the overall temporal patterning of songs and more local spectral adjustments might translate into a general pattern in birdsong evolution [39] [40].

Segregating information types to specific sound dimensions or "channels" might allow the callers to encode this information more accurately [41], [42]. For example, the little auks use complex calls (*classic call* [43]) that carry strong identity signals in both spectral [29] and rhythmic [6] parameters, but contain information on partnership in the spectral and sex in the rhythmic dimension, respectively [44] [6]. For callers that need to signal their identity efficiently to acoustically "mark" their territories, like the tawny pipit, this might mean maximising the level of variability (in this case, the number of recognisable individuals) within one "channel" or dimension.

We could correctly assign songs to their populations of origin, although the precision of this classification was not particularly high. This can be explained by the S-UMAP clustering (Figure 6), which revealed similarities between the two Greek populations and the Spanish site of Rincón de Ademuz. While the similarity of geographically close Greek populations is not surprising, the Spanish site raises some additional questions. Could this be an artifact of excluding other, spectral parameters? Or of the relatively smaller representation of these populations in the dataset?

Tawny pipits are Palaearctic migrants, wintering in the Sahel and moving to breed in their Northern ranges (for this study’s population, Europe [45]). Eastern (Greek) and Western (Spanish) European populations are expected to migrate via Egypt and Gibraltar, respectively, and winter on the opposite sides of Sahel [46], [47], and are thus highly unlikely to meet and mix. One aspect that the Greek and Rincón de Ademuz populations do seem to have in common is an apparently lower local density (CP-G, personal observations during fieldwork). While this might be expected to have some effect on local individuality [48], it remains unclear whether and how local density might influence geographic rhythmic patterns in song.

### Future directions

While we are purely interested in the rhythmic aspect of tawny pipit songs, we suggest that future studies on this and other species take the full picture (i.e., both spectral and a comprehensive set of rhythmic parameters) into account. Note that here, we have decided to use single values or summary statistics for all parameters for each song (so that e.g. integer ratios are presented as minimum, maximum, and mean), following one of standard approaches in bioacoustics. Using the full set of integer ratios of all element pairs might provide a level of detail similar to musical notation and possibly prove useful in further research. The rhythmic signal structure of woodpeckers innate drumming behaviour shows clear phylogenetic differences [49], indicating at gene-driven evolution of its rhythm. How might rhythms develop in vocal learners, such as pipits? A future study, pooling data from more pipit species sampled across some of their distribution, and adding a genetic dimension (as to quantify population divergence and connectivity), should help us disentangle phylogenetic and cultural impacts on song evolution. Whether driven by genes or memes (small cultural units that spread by imitation) [50] [51], if evolution of rhythm occurs, similar patterns might emerge in various populations (or, in fact, be conserved across populations, similarly to human music [52]).

Both individual- and population-scale rhythmic patterns seem to emerge clearly in the species’ song. Rhythmicity seems to emerge as an important aspect of avian communication [53] [5] [54], but the extent to which the animals perceive and use the information carried by this channel remains unclear [55] [56]. Dedicated playback experiments based on artificial songs with modified rhythmic parameters would be of help here. We suggest, however, to first address this issue in a species with more organised, isochronous songs, where these measures are easier to interpret and modify. Some bird species use and spontaneously entrain to rhythmic patterns (e.g. [36] [57] [58]), and songbird vocal athletics (in terms of frequency precision and tempo) are an important part of mating systems in this group [59] [60]. We suggest that parameters such as beat precision might also signal the singer’s quality (similar to how professional musicians produce more exact beats than a layperson).

Not considering the usefulness to birds themselves, this work still provides new and useful insights into their communication systems. Understanding that both identity and population information can be contained within the rhythmic structure can also contribute to improving passive acoustic monitoring of animal populations [9]. This is also a step forward in understanding the origins and function of rhythm, and the evolution of music and communication [20] [21] [22] [23].

### Conclusions

We present the first evidence that modern rhythm analysis methods can reliably describe arrhythmic sequences. The case study of the European tawny pipit population shows that rhythmic differences in such arrhythmic songs can be used to identify both individual singers and their population of origin, the latter previously not identified acoustically with success.

## Methods

### Ethical note

This project was based on already available and previously published acoustic data [61], and as such no specific permits were necessary. The original recordings were collected over the breeding seasons 2015-2022 using non-invasive distance sampling methods (see methodological details in [11]).

### Fieldwork and recordings

Recordings were collected in six European breeding sites of the species (Figure 2). As per the species preference, all these locations were characterised by similar open habitats, with low or limited vegetation, with scattered rock formations, bushes, or trees used as singing posts by the birds [11].

Recordings were made using Sennheiser ME-67 shotgun directional microphones and Marantz PMD 661 recorders, in a focal follow regime with vocal notes to identify the recorded singer.

### Data analysis

#### Data annotation and rhythmic indices

All data were annotated by JO-C in Luscinia (v. 2.24.09.12.02) [13]. Luscinia is a bioacoustics software that allows detailed annotation of vocalisations by manually tracing spectral contours on spectrograms with a brush tool. This approach is particularly useful for sounds with overlapping elements, as it allows the user to separate and measure each element reliably. and ensures precise time measurements, even in songs with substantial overlap between elements or irregular spectral patterns. Thanks to this, we were able to identify song elements of these somewhat chaotic sounds. When needed, background noise was reduced and signal quality improved by adjusting the dereverberation (100%), dereverberation range (50 ms), dynamic range (40 dB), and high-pass threshold (1000 Hz) settings. Every element within a song was then tracked, only including recordings of sufficient quality (i.e. those in which song elements could be clearly distinguished from background noise). All elements were considered, which means that e.g. in the case where long tonal sounds overlapped with a series of short pulses, each element was marked separately.

The start and end times of these elements were then exported from Luscinia and used to calculate a set of temporal and rhythmic indices (Table 2) using a custom made script (based on [29] [5] [4]) in R v. 4.4.2 [62]. Rhythmic indices were calculated for all pairs of element onset times, independently of this overlap.

**Table 2:**
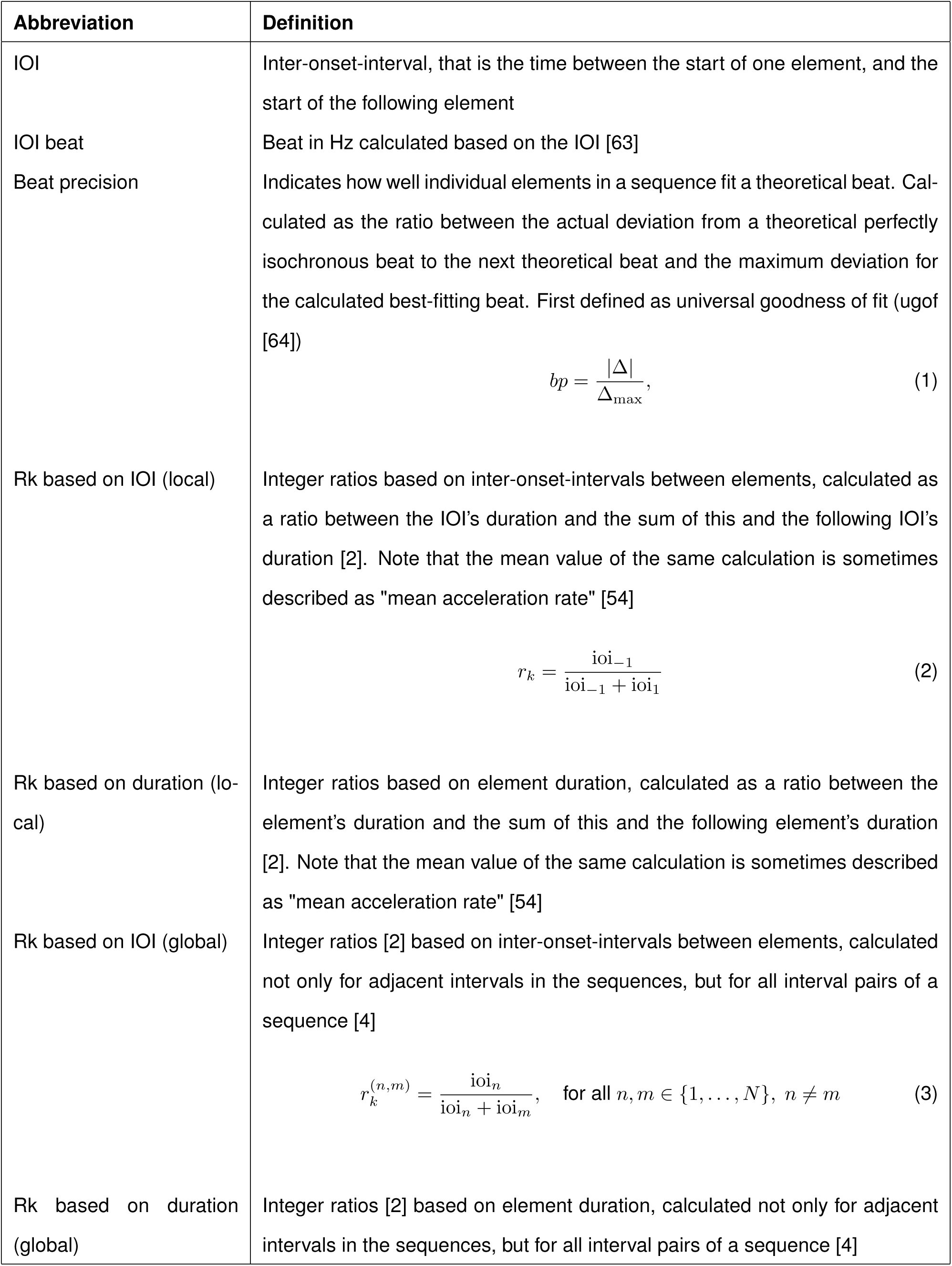
Definitions of the concepts behind rhythmic parameters calculated for each song. For brevity, more commonly used concepts such as durations or number of elements, as well as the specific measures for each parameter (minimum, maximum, mean, coefficient of variation) are not explicitly defined

#### Classification to individual and population

To avoid using underrepresented birds, but maximise the amount of available data, we have only kept songs of birds with at least three recorded songs (1191 songs of 384 individuals in total).

We have then confirmed that the data were appropriate for factor analysis, by conducting the Kaiser-Meyer-Olkin test (KMO function of EFAtools package, [65]). With a KMO score of 0.72, the result was middling and data suitable to conduct a principal components analysis (PCA). We conducted the PCA using prcomp function of the stats package [62], and only retained the PCs with eigenvalues >1 for further analysis (Kaiser’s criterion).

To investigate the *individuality of songs across populations*, we performed a permuted discriminant function analysis (pDFA [14]) pooling all available songs from all individuals and from across the six populations. The pDFA with nested design was conducted using the pDFA.nested function of a script provided by Roger Mundry (based on function lda of the MASS package [66]), using individuals unique ID as a test factor, and population as a restriction factor, and the retained PC scores as input variables.

To investigate the *geographical patterns in rhythmic structure*, we performed a similar pDFA, with population code as the test factor and singers’ ID as a control factor. A Supervised Uniform Manifold Aproximation and Projection (S-UMAP [67]) was created for visual investigation of this clustering using the umap function of uwot package [68], using three components, 200 nearest neighbours (prioritise global over local structure), minimum distance 1 (for tight clusters), and Euclidean metrics.

#### Identity coding

To report cross-species comparable identity metrics [15], we have calculated the Beecher’s information statistic (Hs), which reports a level of individuality carried by an acoustic signal [16]. This was done by first collapsing the raw parameters into a PCA to obtain uncorrelated principal components, and then calculating the Hs based on all obtained principal components (functions calcPCA and calcHs of the IDmeasurer package [69])

To understand the relative importance of each of the rhythmic indices, we calculated the Potential of Individual Coding (PIC [17]) of each parameter using the calcPIC function of the IDmeasurer package [69].

#### Data accessibility

All data and code produced for this work is openly accessible at https://osf.io/h2fpm/overview?view_only=60916d5355de494295a1e9612ca3d40c

## Authors contributions

AB: Investigation. ANO: Conceptualization, methodology, investigation, data curation, writing: original draft, review and editing, project administration, software, visualisation, funding. AP: Methodology, investigation, visualisation, writing: review and editing. CP-G: Investigation, writing: review and editing. JO-C: Methodology, investigation, data curation, writing: review and editing, funding. JT: Investigation, funding, writing: review and editing. LSB: Methodology, software, writing: review and editing, supervision. MP: Investigation. TK: Investigation. TP: Methodology, investigation, writing: review and editing, funding. VB: Investigation, funding.

## Acknowledgements

The authors would like to thank Roger Mundry for sharing his pDFA code, Marianne Sarfati for her assistance with the Luscinia software and Pavel Linhart for his valuable insights on identity metrics.

